# An olfactogenetic approach identifies olfactory neurons and brain centers directing negative oviposition decisions in *Drosophila*

**DOI:** 10.1101/206342

**Authors:** Sonia G. Chin, Sarah E. Maguire, Paavo Huoviala, Gregory S.X.E. Jefferis, Christopher J. Potter

## Abstract

The sense of smell influences behaviors in animals, yet how odors are represented in the brain remains unclear. The nose contains different types of olfactory sensory neurons (OSNs), each expressing a particular odorant receptor, and OSNs expressing the same receptors converge their axons on a brain region called a glomerulus. In *Drosophila*, second order neurons (projection neurons) typically innervate a single glomerulus and send stereotyped axonal projections to the lateral horn. One of the greatest challenges to studying olfaction is the lack of methods allowing activation of specific types of olfactory neurons in an ethologically relevant setting. Most odorants activate many olfactory neurons, and many olfactory neurons are activated by a variety of odorants. As such, it is difficult to identify if individual types of olfactory neurons directly influence a behavior. To address this, we developed a genetic method in *Drosophila* called olfactogenetics in which a narrowly tuned odorant receptor, Or56a, is ectopically expressed in different olfactory neuron types. Stimulation with geosmin (the only known Or56a ligand), in an *Or56a* mutant background leads to specific activation of only the target olfactory neuron type. We used this approach to identify which types of olfactory neurons can directly guide oviposition decisions. We identified 5 OSN-types (Or71a, Or47b, Or49a, Or67b, and Or7a) that, when activated alone, suppress oviposition. Projection neurons partnering with these OSNs share a region of innervation in the lateral horn, suggesting that oviposition site-selection might be encoded in this brain region.

**Significance Statement:** The sense of smell begins by activation of olfactory neurons in the nose. These neurons express an olfactory receptor that binds odorants (volatile chemicals). How the sense of smell is encoded in the brain remains unclear. A key challenge is due to the nature of olfactory receptors themselves - most respond to a wide range of odorants - so it is often impossible to activate just a single olfactory neuron type. We describe here a novel approach in *Drosophila* called ‘olfactogenetics’ which allows the specific experimental activation of any desired olfactory neuron. We use olfactogenetics to identify olfactory neurons and brain regions that guide egg-laying site selection. Olfactogenetics could be a valuable method to link olfactory neuron activities with circuits and behaviors.

## Introduction

The sense of smell is an ancient and vital sensory system but how olfactory information is processed in an animal’s brain remains unclear. Organisms ranging from insects to humans use the olfactory system to detect volatile chemicals (odorants). These odorants act as environmental sensory cues representative of the physical objects that emit them. Upon detection of these volatile chemicals, the olfactory system must detect, interpret, and then guide behavioral actions based upon these cues. Olfactory perception results from the activation of individual olfactory sensory neuron (OSN) types found in olfactory organs (such as the nose), which each express specific odorant receptors responsible for detecting a near-infinite variety of odorants. The number of OSN types, defined as expressing a specific olfactory receptor, differs among species, with humans having 388 OSN types, mice having 1200 OSN types, and vinegar flies having 62 OSN types (1, 2). Axons of OSNs that express the same olfactory receptor project directly to a specific region of the brain called a glomerulus, which is the primary organizing unit for the olfactory system. The collection of all glomeruli is called the olfactory bulb in mice, and the antennal lobe in insects. Dendrites of second order neurons (mitral/tufted cells in mammals, and projection neurons in insects) typically innervate a single glomerulus. Thus a glomerulus acts as an organizing center to match OSNs to their cognate second order projection neuron partners. Projection neurons then send axons to the olfactory cortex (piriform cortex in mammals, lateral horn in insects) that are centers for further olfactory processing. How does activation of individual OSN types contribute to a complex behavior? This has been difficult to study because most natural odors contain complex mixtures of odorants, and most odorants can activate several different olfactory receptor classes at varying degrees (3). This aspect of the olfactory system makes a precise odor-driven experimental study of OSNs extremely challenging. A goal of this study was to develop genetic tools in an animal model that could be used to investigate the contributions of individual classes of OSNs to odor-guided behaviors.

The vinegar fly, *Drosophila melanogaster*, is a powerful genetic model for investigating sensory perception. As a highly olfaction-driven organism, the fly uses its sense of smell to direct all essential behaviors from locating food, navigating space, mating with the correct species, and finding locations to lay eggs (4). Many of these olfactory behaviors occur on highly odiferous rotting fruits, which serve as both food source and oviposition substrate. The fly's olfactory system must filter through a complex sensory world in order to obtain ethologically relevant odor information and behave appropriately. Comprehensive screens have matched the identity of each odorant receptor type to the OSNs that express them, and the response profiles of many OSNs have been determined (3, 5, 6). Furthermore, the glomerular targets for each OSN type in flies has been genetically mapped (7, 8) - the most comprehensive olfactory map for any organism. Like the mammalian olfactory system, *Drosophila* OSNs expressing the same odorant receptor all target a single glomerulus. However, unlike in mammals, insect OSN targeting does not depend on which odorant receptor is expressed, and anatomical locations of individual glomeruli in the antennal lobe are highly stereotyped. The projection neurons innervating many glomeruli have also been genetically identified and are also highly stereotyped in their anatomical targeting patterns in the lateral horn (9, 10). Based on these axonal innervation patterns into the lateral horn, it appears that projection neurons re-organize olfactory information from the antennal lobe to the lateral horn such that the different regions of the lateral horn represent biologically relevant stimuli- such as food versus pheromone odors, and possibly aversive odors (9). The identification of additional lateral horn domains has been hampered by the previously mentioned experimental challenge in linking individual OSN activities to discrete olfactory behaviors.

A critical behavior for species survival in most insects is determining where to lay eggs (oviposition) (11). Females that choose high quality egg-laying substrates ensure the health of both eggs and the resulting larvae. Since larvae cannot fly and usually remain close to the oviposition site, their ability to survive depends heavily on the patch of food on which they hatch. Thus, a mated female fly must balance her own nutritional and safety needs with the needs of her future offspring to maximize nutrient intake and protection from parasites and disease-causing microorganisms (12). Many sensory cues guide oviposition decisions: taste (gustation), texture, vision, and olfaction (13, 14). The chemical senses, in particular, strongly contribute to oviposition decision-making. Several individual olfactory receptors have indeed been associated with oviposition decisions (15–20). Nonetheless, how the brain processes olfactory information to generate egg-laying behavior is not well understood.

To study how defined OSN types give rise to olfactory signals used by the brain to guide oviposition decisions, we generated a novel genetic tool that allows highly specific odorant-directed activation of individual OSN classes. We call this method ‘olfactogenetics’ to reflect that it uses a volatile odorant to activate genetically defined types of olfactory neurons. This approach offers several advantages over existing genetic activation methods (such as optogenetics (21) and thermogenetics (22)); for example, induced olfactory activities will more closely mimic natural odorant stimulations and do not rely on sophisticated equipment designs. We used the olfactogenetic method to systematically identify OSNs that, when activated, mediate oviposition choices. By linking the glomerular targets of the OSNs to likely activated projection neurons, we identified a region of the lateral horn that might represent negative oviposition cues.

## Results

### An olfactogenetic method to activate specific OSN types

We developed a novel olfactogenetic method for specific OSN activation that takes advantage of an unusually specific odorant-receptor to odorant pair identified in *Drosophila* (16). In a screen of olfactory sensilla, the odorant geosmin was found to activate only a single olfactory neuron class (antennal basiconic 4B) expressing the Or56a receptor (16). This suggests that Or56a is the only odorant receptor that is activated by geosmin, and deleting the Or56a receptor should result in a fly that is completely anosmic to geosmin. Furthermore, ectopically expressing Or56a in a different OSN type should then confer geosmin-induced responses to that olfactory neuron. The olfactogenetic approach thus requires three components: [1] Or56a mutant (*Or56a*^-/-^) animals to eliminate wild-type responses to geosmin, [2] a *UAS-Or56a* transgene to drive *Or56a* in olfactory neurons, and [3] *OrX-GAL4* lines to direct which types of olfactory neurons express *UAS-Or56a* (**Fig 1A**). We generated *Or56a*^-/-^ animals and confirmed by single sensillum recordings (SSR) that responses to geosmin in the ab4 sensillum were abolished (**Fig 1B, Fig S1**). To verify the identity of the *Or56a*^-/-^ mutant sensilla, we stimulated the sister olfactory neuron (ab4A) that expresses Or7a with the activating pheromone 9-tricosene (15), and found that ab4A neurons fired normally. This confirmed that in the *Or56a*^-/-^ mutant sensilla, only the large spiking amplitude A neuron fires (Or7a) while the B neuron (Or56a) is completely silenced indicating the fly no longer expresses an olfactory receptor capable of detecting geosmin. Consistent with a previous study examining geosmin responses for all olfactory neurons in *Drosophila* (16), we found ab4A (Or56a+) neurons were the only olfactory neurons activated by geosmin (see behavior results below; data not shown).

**Figure 1.**
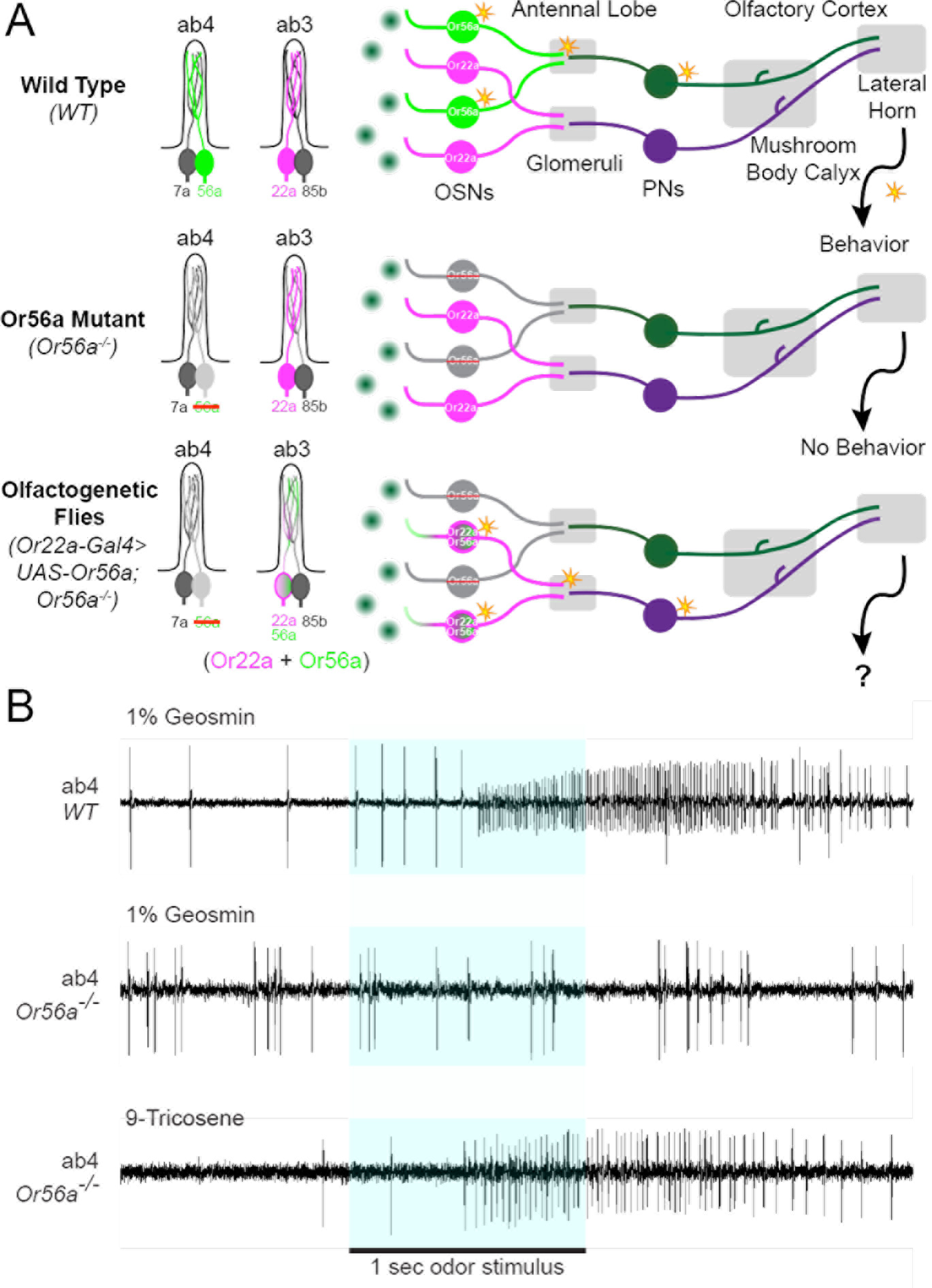
Or56a-geosmin olfactogenetic method for investigating odor-guided behaviors. **A)** Schematic of olfactogenetic approach. In Wild Type (WT) conditions, the ab4 sensillum contains 2 OSNs : the ‘A’ neuron expresses Or7a (grey) and the ‘B’ neuron expresses Or56a (green). The ab3 sensillum also contains 2 OSNs: the ab3A neuron expresses Or22a (magenta) and the ab3B neuron expresses Or85b (grey). Geosmin (dark green circles) activates (orange star) only Or56a-positive neuronsand not other neurons. Mutating the *Or56a* receptor results in an olfactory system that does not detect nor behaviorally respond to geosmin. To create olfactogenetic flies, we use the GAL4/UAS system to ectopically express *UAS-Or56a* in a specific olfactory neuron (*e.g., Or22a-GAL4*, magenta plus green with magenta outline) in an *Or56a*^-/-^ mutant background. This allows the odorant geosmin to activate olfactory neurons with high specificity to drive olfactory behaviors. **B)** Single sensillum recordings (SSR) of ab4 sensilla. The ‘B’ neuron in an *Or56a*^-/-^ mutant no longer responds to geosmin. The ‘A’ neuron in an *Or56a*^-/-^ mutant responds normally to 9-tricosene. ab, antennal basiconic. PNs, Projection Neurons. OSNs, Olfactory Sensory Neurons.

### Ectopically expressed Or56a activates olfactory sensory neurons through stimulation with geosmin

Four types of sensillum, classified according to morphology, cover the fly’s olfactory organs: basiconic, intermediate, trichoid, and coeloconics (6, 23). The olfactory neurons in these sensilla also tend to respond to different types of odorants: trichoid olfactory neurons respond to pheromones, intermediate olfactory neurons respond to kairomones (odorants from other species), basiconic olfactory neurons respond to a diverse set of environmental odorants, and coeloconic olfactory neurons respond to amines and acids (3, 23). Previous studies indicate that the intracellular molecular environment of different sensillum types may differ. For example, signaling in antennal trichoid (at) 1 has been shown to be enhanced by the presence of the odorant binding protein LUSH and sensory neuron membrane protein (SNMP1) (24, 25), and a recent study de-orphanizing Or83c in antennal intermediate (ai) 2 suggests intermediate sensilla may possess intra-sensillar components more similar to trichoids than basiconics (26). Or56a normally expresses in the antennal basiconic sensillum ab4 so it was important to validate that ectopic expression of this receptor in non-native sensillum types produces a physiological response. Thus, we used SSR to determine if ectopically expressed Or56a in non-ab4B neurons would be functional and activated by geosmin.

We sampled the olfactogenetic activation of neurons from both olfactory organs (antenna and palp) and three of the four major sensillum types. *UAS-Or56a* was expressed in ab3A (*Or22a-GAL4*), palp basiconic (pb) 1B (*Or71a-GAL4*), ai2B (*Or23a-GAL4*), at1 (*Or67d-GAL4*) and at4A (*Or47b-GAL4*). Since Or56a is natively expressed in a basiconic sensillum (ab4), it was expected that ectopic expression of Or56a in other basiconic sensilla would be able to confer a geosmin response. This was the case in both the antenna (ab3A) and maxillary palp (pb1B) (**Fig 2**). While primarily testing the response of ab3A (*Or22a-GAL4*>*UAS-Or56a*), we observed that geosmin stimulation also appeared to cause a change in firing rate of its neighboring B neuron (**Fig S2**). The ab3B neuron responded to the pure mineral oil control. Since neurons housed in the same sensillum share hemolymph, the firing of one neuron has the ability to inhibit firing in its neighbors through the phenomenon of ephaptic coupling (27). Ephaptic coupling along with ab3B’s native response to the vehicle mineral oil likely explains why presentation of geosmin to activate ab3A also silences ab3B.

**Figure 3.**
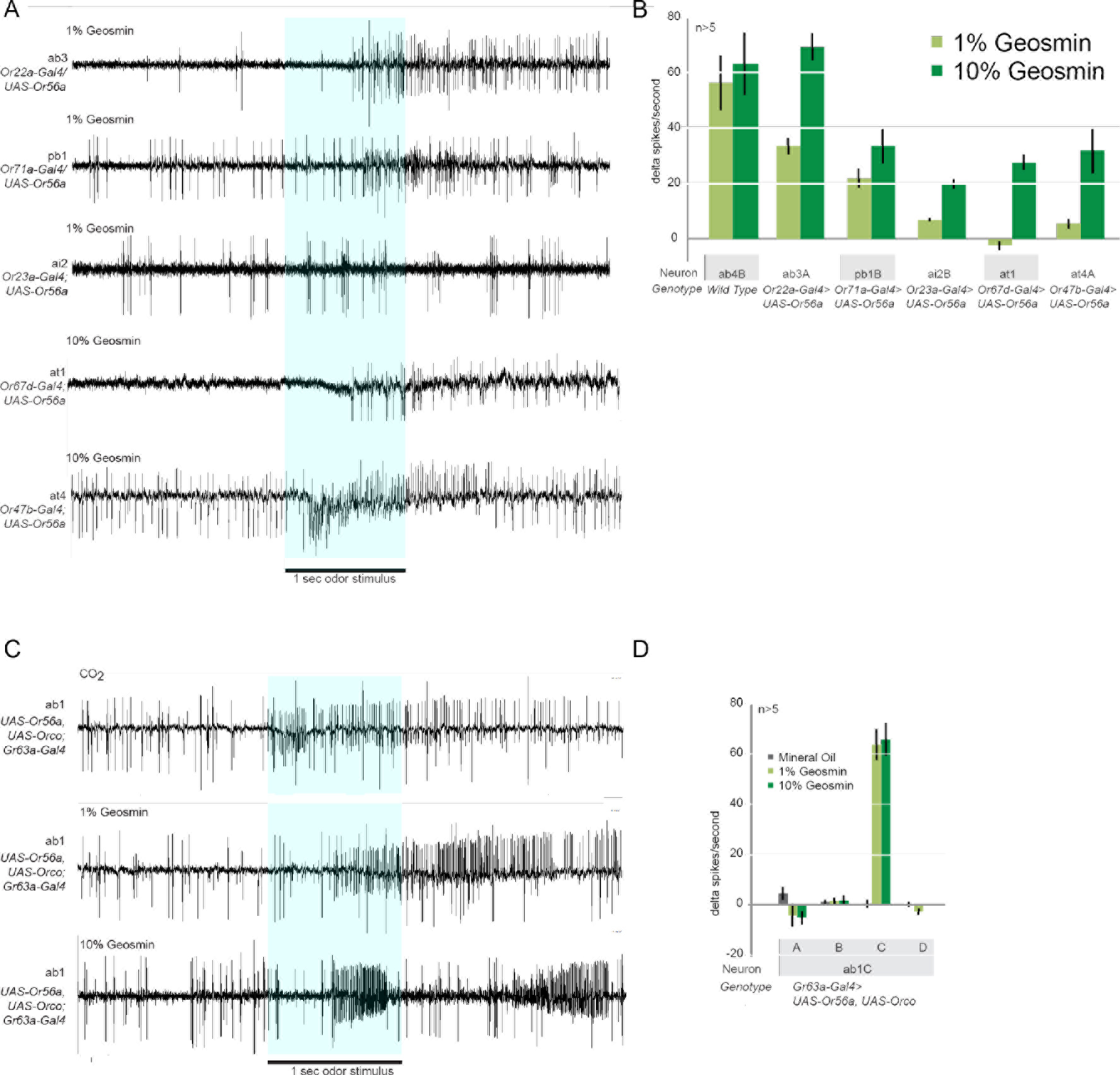
Ectopic expression of *Or56a* in olfactory neurons confer response to geosmin. **A)** Representative single sensillum recordings of basiconic, intermediate, and trichoid sensilla containing neurons expressing Or56a. The blue bar highlights the 1-second odor pulse. **B)** Quantification of recordings. **C)** Representative SSR of ab1 sensilla that contains 4 olfactory neurons. The ‘C’ neuron expresses Gr21a/Gr63a and is sensitive to CO_2_. **D)** Quantification of recordings. Error bars are SEM.

The intermediate sensillum neuron ai2B also reliably increases firing, but only by <10 Δspikes/second (**Fig 2**). It is possible that non-basiconic intermediate sensilla contain different molecular components (e.g., odorant binding proteins, co-receptors, or odorant degrading enzymes), which might be responsible for weak activation. Therefore, we also tested expressing *Or56a* in ai2A (*Or83c-GAL4*), the companion neuron to ai2B in the same sensillum (**Fig S2**). The ai2A neuron was robustly activated by geosmin (30-50 Δspikes/sec), suggesting that basiconic Ors can function normally in intermediate sensilla. Expression of Or56a in trichoid sensillum neurons at1 and at4A conferred activation by geosmin, but only at the higher (10%) geosmin concentration (**Fig 2A,B**).

This suggests trichoid sensilla, like intermediate sensilla, may contain different molecular components that normally optimize responses to pheromones, but may reduce the responses to non-pheromone odors (24, 25). Nonetheless, the level of olfactogenetic activity induced in intermediate and trichoid sensilla should be sufficient to drive olfactory behaviors (28).

Olfactory sensilla most commonly express canonical Ors along with the coreceptor Orco (29). However, the ab1C neuron expresses two gustatory receptors - Gr21a and Gr63a - that together detect CO_2_ (30). The Or56a-geosmin olfactogenetic tool could potentially be useful in studying gustatory receptor neuron (GRN) functionality. GRNs do not express Ors or Orco (31, 32). Given that the mechanism of activation and necessary intracellular signaling components of Grs are not well defined, it was unclear whether an Or could be used to activate a GRN. Expressing both *Or56a* and *Orco* in ab1C (*Gr63a-Gal4*>*UAS-Or56a*, *UAS-Orco*) and stimulating with geosmin indicates that ab1C can be robustly activated using Or56a-geosmin olfactogenetics (**Fig 2C, Fig 2D**). Notably, 1% Geosmin stimulation to ab1C elicited typical firing dynamics with a continuous burst of firing after odor presentation that gradually returned to baseline. Interestingly, this was not the case with 10% Geosmin stimulation. In 5 of the 6 recordings using 10% geosmin, we observed a biphasic response where, upon initial presentation of the odor, the ab1C neuron fired rapidly followed by a sudden decrement to no firing, and then a second burst of firing erupted shortly thereafter (**Fig 2C**). This exclusively occurred in response to high concentration geosmin stimulation. While Ors likely make a complex with Orco to form ion channels, several studies have shown that second messengers are necessary for normal responses to odorants (33, 34). The difference in firing dynamics in ab1C suggests that Grs may have different downstream second messengers that act at a different time scale than those of Ors, or perhaps that Gr-specific cellular machinery can act to temporarily repress Orco-Or56a receptor complex signaling.

### Two-choice oviposition assay relies on olfactory signaling

Many sensory cues, including olfactory cues, influence oviposition choice in female flies. We established an oviposition choice assay in which only olfaction was used as a differentiating cue. In this assay, three wells in a dissection spot glass were filled with odorless 1% agarose to serve as a suitable egg-laying substrate. Geosmin was added to the agarose in one of the three wells. Egg-laying behaviors were conducted in a completely dark, humidified incubator for 23 hours. An oviposition index (OI) was calculated as the number of eggs laid in the odor well minus the average number of eggs laid in the other two wells, divided by the total number of eggs; an OI =1 indicates all eggs were laid in the odor well, whereas an OI < 0 indicates flies avoided laying eggs in the odor-well. Since female flies could touch the agarose gel containing geosmin, it remained possible that chemosensory receptors on the feet or ovipositor might contribute taste cues and affect oviposition in this olfactory assay. To rule this out, we performed experiments with olfactory mutant flies. We tested three genotypes: 1) the *Or56a* mutant 2) the *Orco* mutant (which disrupts all Or-type signaling) and 3) flies mutant for ionotropic receptor co-receptors (*Ir8a* and *Ir25a*), *Orco*, and *Gr63a* (which disrupts almost all olfactory signaling). All three olfactory mutant genotypes exhibited neutral oviposition indices to geosmin, indicating that knockout of *Or56a* eliminates behavioral response to geosmin in our oviposition assay, geosmin is likely not detected by other chemosensory neurons, and gustatory signaling does not contribute to the behavioral response in this assay (**Fig 3B**). Changing the position of the geosmin odorant well did not change oviposition scores, and flies showed equal exploring for all wells, suggesting no edge effects in this assay (**Fig S3**).

**Figure 3.**
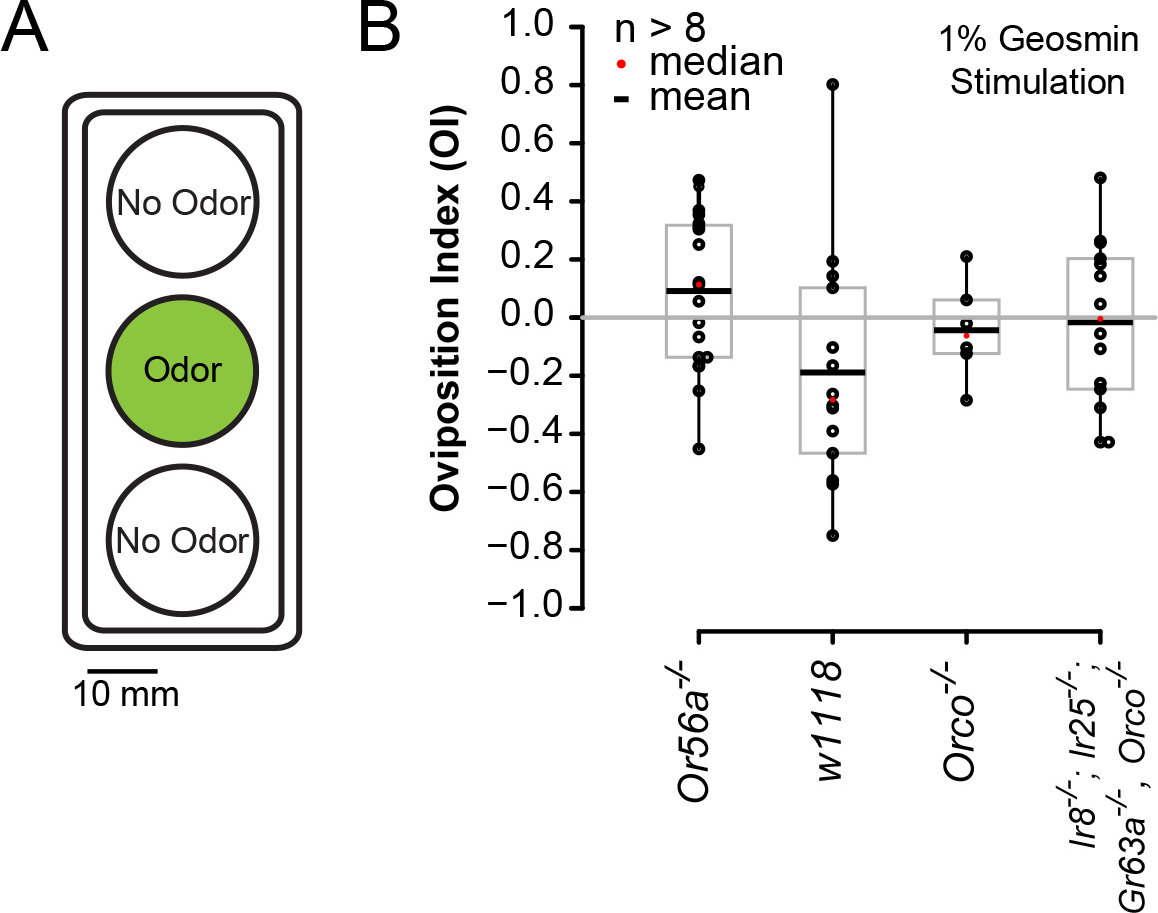
Behavioral effects in the two-choice oviposition assay using geosmin rely on olfaction. **A)** Schematic of oviposition assay. **B)** Olfactory mutant animals do not exhibit behavioral responses to geosmin. Control *w*^*1118*^ animals are slightly repelled by geosmin. The oviposition index is calculated as the # of eggs laid in odor well - the average number eggs laid in the no-odor wells, divided by the total number of eggs. The different genotypes are not statistically significant as determined by a Dunnett’s Many-to-One Comparisons Test.

### Single OSN-types mediate negative oviposition decisions

We systematically performed an olfactogenetic screen using twenty-three *OrX-Gal4* lines to identify types of olfactory neurons that, when activated, contributed to oviposition choices in females. These *OrX-GAL4* lines were chosen because they drive expression in OSNs with target glomeruli for which partnering PN morphology is known. We reasoned this might help elucidate downstream olfactory circuit signaling. The results of the olfactogenetic screen are shown in Figure 4. Our data indicate that activating just a single OSN type is, indeed, sufficient to elicit a reproducible oviposition behavior. A range of oviposition indices was observed among the different *OrX-GAL4* lines used. Notably, all statistically significant *OrX-GAL4* responses from the screen demonstrated negative oviposition decisions (**Fig 4A**). Many of the OrX-expressing olfactory neurons (*e.g*., Or71a, Or49a, Or7a) have been previously shown to be involved in oviposition decisions (**Supplemental Table 1**). To determine if the olfactogenetic approach might recapitulate the behavioral responses of native odorant-to-Or responses, we examined the responses of three odorants (4-ethylguiacol, 9-tricosene, methyl laurate) that are fairly specifically tuned towards their respective odorant receptors (Or71a, Or7a, Or47b) (**Fig 4B**). The behaviors mediated by the olfactogenetic approach mimicked the oviposition odorant-induced responses towards 4-ethyl guiacol and 9-tricosene, but not towards methyl laurate. The lack of recapitulation in the methyl laurate case may be due to the activation of other chemosensory neurons that simultaneously respond to methyl laurate (35, 36) that could possibly modify oviposition behavioral decisions. Altogether, this highlights how an olfactogenetic approach can be used to investigate which behaviors are solely mediated by the targeted olfactory neurons.

**Figure 4.**
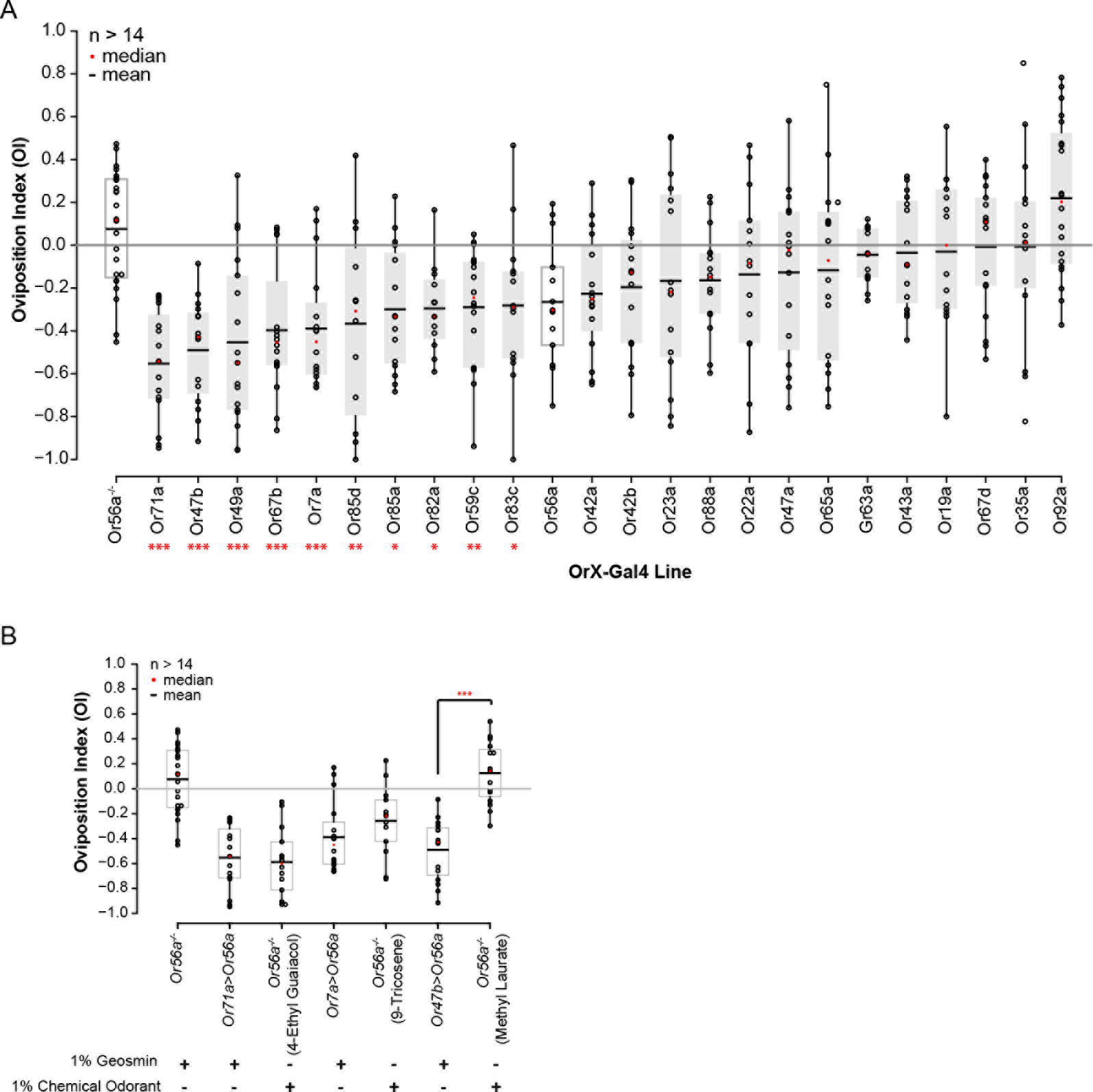
Olfactogenetic activation of specific OSN types mediates negative oviposition. **A)** Oviposition assays using geosmin. *OrX-Gal4* lines were combined with *UAS-Or56a* in the *Or56a*^-/-^ mutant background (gray bars). *Gr63a-GAL4* combined with *UAS-Or56a and UAS-Orco* in the *Or56a* background (gray bar). Mutant *Or56*-/- and wild-type (*Or56a*) responses denoted by white bars. **B)** Oviposition assays comparing results obtained from an olfactogenetic approach to those obtained using the indicated odorants. Statistics are a Dunnett’s Many-to-One Comparisons Test compared to *Or56a*^-/-^;, Stars indicate p-values: ‘*’<0.1, ‘**’ <0.05, ‘***’ < 0.001.37

### A region of the lateral horn may mediate negative oviposition decisions

The olfactogenetic screen took an unbiased approach towards identifying *OrX-Gal4* lines that could direct oviposition decisions. We next examined the projection patterns for the PNs predicted to be the primary signaling partners for each of the identified OSNs (Fig 5A). Interestingly, the PNs predicted to guide negative oviposition decisions were found to be significantly more similar to one another than by chance based on their morphology in the lateral horn (**Fig 5B, Fig S4**). Indeed the axonal arbors of each negative oviposition PN inhabited an anterior-central region of the lateral horn (**Fig 5B**). Closer examination of these PN traces indicated that for VC2 and DL5, only the medial branches, rather than whole axonal pattern, shared morphological similarity to the other negative oviposition neurons. This implies that structurally similar axonal segments may define functional domains in the lateral horn and that different branches of an axon from a PN, by potentially connecting with distinct postsynaptic partners, might encode functionally different olfactory information.

**Figure 5.**
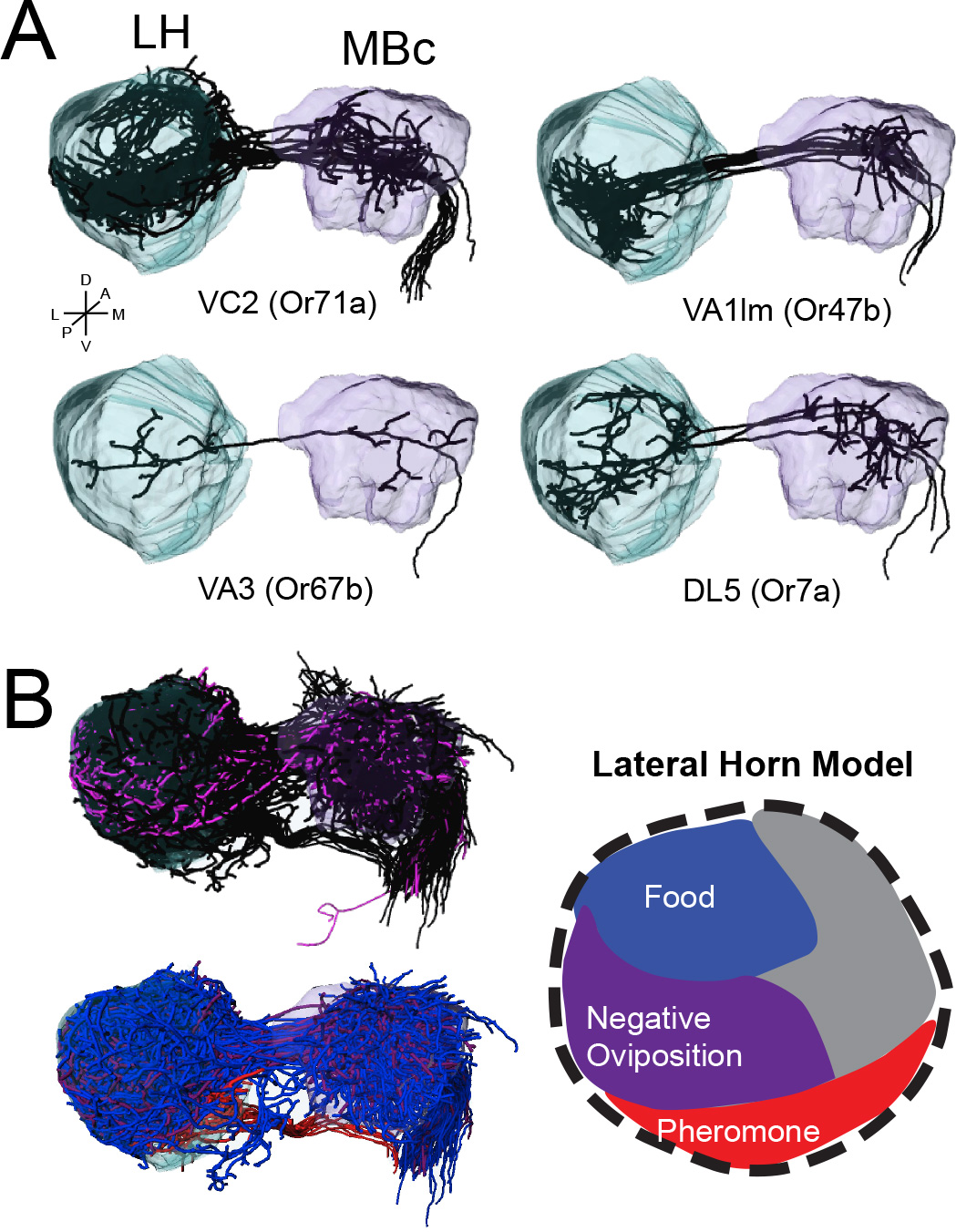
Representation of negative oviposition olfactory cues in the Lateral Horn. **A)**Projection neuron traces corresponding to the listed OSN-type for the most statistically significant responses in the olfactogenetic oviposition assay (p>0.001). LH, Lateral Horn; MBc, Mushroom Body calyx; A, anterior; P, posterior; D, dorsal; V, ventral; L, lateral; M, medial. **B)** Comparison of negative oviposition PNs (purple) to all other PN types (black in top trace, blue and red in categorized bottom).

## DISCUSSION

Chemosensation is considered one of the most primal senses as all animals (from bacteria to humans) use chemical information to interact with their environments. The olfactory system has the ability to detect volatile odorants that drives integral survival behaviors such as finding nutritious food, identifying an attractive mate, avoiding ingestion of disease-causing microbes and toxins, and influencing oviposition, courtship, aggregation, flight, and aggressive behaviors (17, 37–39). The chemical world contains a large diverse number of compounds. How does the brain make sense of it all?

### A genetic olfactogenetic approach for the dissection of olfactory behaviors

The contribution of olfactory neuron types in guiding olfactory signaling has been technically challenging to investigate. In contrast to other sensory systems, such as audition or vision, which benefit from the ability to precisely control the experimental sensory input, the olfactory system is not as amenable to such systematic experimental investigations. This is because most olfactory neurons do not respond to a single odorant, and most odorants activate many olfactory neuron types, making it difficult to link which olfactory neuron activities in response to an odor are actually the main drivers for the resulting olfaction-guided behavior. To overcome this hurdle, we took advantage of a usually specific odorant-odorant receptor pair (geosmin/Or56a), and developed an olfactogenetic approach that uses a natural odorant stimulus to activate an experimentally defined olfactory neuron population. This allows discrete olfactory neuron types to be activated in a more natural manner, such as towards odor plumes or in response to odor gradients. Geosmin is an odor that has the dynamic features of natural olfactory stimuli. This eliminates confounding factors in the interpretation of behavior often encountered by other experimental methods aimed at neuronal activation like optogenetics (which can lead to phototaxis) and thermogenetics (which can lead to thermotaxis). Being able to use the same odorant to compare results among behavioral assays also helps to eliminate effects of varying volatility between different odorants, and the system allows for the study of receptor neurons whose receptors have no known activating ligands.

In initial studies of OSNs and their firing rates, relatively high concentrations of odorants (1%) were used to elicit olfactory neuron firing. For example, in one of the first instances where OSNs were systematically screened to a panel of odorants (3, 40), responses were categorized as ‘hits’ if responses were greater than 50 spikes/second. Responses of ~150-200 spikes/second were considered as reflecting ‘real’ odorant-to-Or matches. A recent study (28) used an elaborate and sophisticated optogenetic setup to tightly control stimulus intensity towards individual moving flies. The study showed that high levels of OSN activity may not be required for generating behavior. In some cases, lower induced activity of the olfactory neuron of around 40-50 spikes/second generated stronger behavior. This supports the efficient coding hypothesis (41) that postulates the level of a stimulus should match the level of neuronal firing in natural environments where an animal has evolved to survive. This optimizes the neuron’s metabolic consumption and dynamic range. While, to our knowledge, extensive studies have not been conducted to quantify concentrations of natural odorants, natural odorants rarely come in the extremely high concentrations used in lab studies. Therefore, more likely than not, sparse coding is used in sensory systems, and weak activation of sensory neurons are significant to the animal’s perception of its environment. The Or56a-geosmin olfactogenetic approach leads to reproducible (20-60 spikes/sec) increases in olfactory neuron signaling, thus likely reflecting activation of an olfactory neuron to ethologically relevant odorant concentrations. Interestingly, it was difficult to over-activate an olfactory neuron using this approach: maximal olfactory neuron responses plateaued at ~60 spikes/sec even when geosmin levels were increased to 10% or more. This was an unexpected, yet advantageous, aspect of the olfactogenetics approach as experimental over-activation of an olfactory neuron by other genetic methods often leads to inhibition (23), which could confuse behavioral interpretations.

### Oviposition Decisions are complex sensory choices

*Drosophila* lay eggs on their food substrate- rotting fruit- so chemosensation plays a large role in oviposition choice since smell and taste provide essential information about the composition of a food source (such as nutritional content and toxicity). Thus far, the primary studies on chemosensation in oviposition have involved Grs, which are found in many body regions that come in contact with food sites, such as the labellum, legs, and ovipositors. While flies are generally attracted to calorie-rich sugar substrates and avoid substrates that contain bitter compounds, interestingly, oviposition sites can change based on the context of the decision. Laying eggs on a bitter substrate may confer survival benefits in the form of deterring parasitic predators or protecting eggs from fungal or microbial infections. However, on larger and/or physically distant patches, larval foraging costs would be high, necessitating large energy expenditure in order to reach a sugar patch to eat. Therefore, under these conditions, it is more advantageous for the female fly to directly lay eggs on sweet, nutrient-rich substrates.

Olfaction also plays an important role in oviposition choices. However, it can often be difficult to distinguish whether a chemical functions solely as an odorant, or also as a tastant. For example, a bitter volatile chemical could potentially be smelled by the olfactory system as well as tasted by the gustatory system. This seemingly semantic distinction is important to make because it appears that receptors that detect the same chemical on different body regions can mediate opposing behaviors. This is thought to be true in the case of bitter compounds eliciting different behavioral valences in oviposition. Gr66a, a bitter receptor that detects a compound commonly used in egg laying assays called lobeline, causes aversion when activated on the legs but egg-laying attraction when activated in the labellum (42). A similar phenomenon has been observed with olfactory vs. gustatory responses to acetic acid (43). The integration of these two sensory modalities along with elements of the egg-laying environment such as patchiness of food resources illustrates that oviposition choice is a complex decision making task.

### Oviposition decisions based on single olfactory neuron activities

Five olfactory receptors have been specifically associated with oviposition. Or19a and Or49a are implicated in avoidance of larval parasitization by wasps (17, 19). Or19a mediates positive oviposition and responds to citrus volatiles repellent to wasps, and Or49a detects parasitoid wasp semiochemicals which female flies want to avoid during oviposition. Or56a and Or71a have been implicated in avoiding the negative effects of infection by microorganisms (16, 44). Or56a detects geosmin, which is emitted by harmful microorganisms (16, 45, 46), and Or71a promotes attractive oviposition because it is thought to detect antioxidants in food that can attenuate oxidative stress resulting from exposure to toxins (47, 48). Finally, Or7a has been shown to detect the social pheromone 9-tricosene and mediates geographical tagging of food sites by males used to attract females (15). 9-tricosene has also been shown to positively stimulate oviposition through Or7a (15).

In order to systematically screen for and identify more olfactory inputs involved in female oviposition, we used the olfactogenetic approach to test twenty-three olfactory receptor Gal4 (*OrX-GAL4*) lines (**Supplementary Table 2**). The top five statistically significant hits (p < 0.001) correspond to neurons expressing Or71a, Or47b, Or49a, Or67b, and Or7a. The major commonality among these receptors is that they detect social chemical cues, either from the same species, or as hallmarks of other insects. Or47b and Or7a have both been shown to specifically respond to pheromones that male and female flies can use to influence individuals of the opposite sex, and Or49a is activated by chemicals that parasitic wasps deposit on substrates that they have visited (15, 19, 35). In the context of egg laying, the olfactogenetic results could indicate that cues such as density of both conspecifics and interspecifics, and the presence or absence of parasites that infect larvae, are most relevant to a female fly’s oviposition decisions.

Interestingly, we only saw statistically significant negative oviposition behavior. Stimulating Or92a OSNs, neurons that contribute to attraction to apple cider vinegar (49), is the only behavioral result that yielded a positive average oviposition index, but this result was not statistically significant, and while Or71a, Or19a, and Or7a have been behaviorally shown to detect positive oviposition cues, our OSN activation screen produced no attractive oviposition when we olfactogenetically stimulated these classes of OSNs. This can be explained in several ways. First, the parameters of behavior assays, especially oviposition, can influence behavioral results. The assays used to identify Or71a, Or19a, Or49a, and Or56a as mediators of oviposition behavior were performed under conditions where odorants were presented with fly food. The chemical components of a naturalistic odor such as fly food can interact with each other in unpredictable and complicated ways. Insect studies show that background odor can indeed change behavior and physiology of olfactory neurons (27, 50, 51). We hypothesize that since our oviposition assay is agarose-based rather than food-based, we are likely minimizing the olfactory background and experimentally only getting low-level activation of single classes of OSNs that project to a single glomerulus. As such, the olfactogenetic screen may lead to the identification of those olfactory neuron classes that are sufficient to drive behaviors on their own. As an extension of this, an olfactory response that needed a specific combination of olfactory neuron activities would not be picked up in the screen. For oviposition, single olfactory neuron classes only had ‘negative’ valences. This suggests that the major contributions for single olfactory neuron classes regarding oviposition may be towards avoidance, and it is possible that attraction requires the activation of several olfactory inputs and glomeruli rather than a single glomerulus.

### Identification of a negative oviposition region in the Lateral Horn

Our results support previous findings that the lateral horn functions as a categorizer of salient olfactory information. Previous studies defined lateral horn domains based on the entire axonal morphology of the PNs (9). Our analysis of PNs involved in negative oviposition suggests that information may be organized based on axonal subsegments. For each PN predicted to guide negative oviposition behaviors, the PN axons shared a dorsal posterior segment. The non-oviposition anterior branch of the cup-shaped PN neuronal target regions may confer an as yet unknown, yet shared, biological significance
in the lateral horn as they localize together.

### Using olfactogenetics to investigate odor coding

There are many hypotheses about how the brain processes incoming olfactory information. The ‘labeled line’ hypothesis supported by studies identifying dedicated Ors reacting to highly specific odorants operates under the assumption that highly biologically relevant stimuli are encoded as labeled lines of information. It is postulated that most receptors will have a ‘most relevant’ ligand yet to be identified (52). This extreme seems unlikely as the olfactory world of an animal, like the vinegar fly, contains more important biologically relevant stimuli that it needs to respond to than olfactory neuron types. As evidenced by contradicting behaviors seen in different assays (38, 53), even olfactory circuits previously thought to be labeled lines for attraction or repulsion do not absolutely produce the same behavior in all contexts. The ‘combinatorial code’ hypothesis stipulates that odorant information is processed and acted upon by the combinatorial activity of many olfactory neurons (54). The antennal lobe acts to linearly summate all inputs from activated and inhibited glomeruli and/or use coincidence detection of simultaneously activated OSNs to determine odor identity and direct behavioral responses (28, 55, 56).

The olfactogenetics approach allows rigorous experimental testing of the labeled line hypothesis by enabling each olfactory neuron type to be activated and assayed for behaviors directed by activity of only that olfactory neuron. It might help to distinguish between behavioral situations guided by labeled lines (like negative oviposition) from those that require combinatorial signaling to drive behaviors. It is also possible that labeled lines exist primarily as modulators of combinatorial signaling. Combinatorial signaling may implicate the behavioral context of the each odorant with labeled lines modulating the overall response to each particular situation. Further study using the olfactogenetics approach could identify still more olfactory neuron types involved with imparting important olfactory information to strongly modulated olfactory circuits or influencing the activity of other OSNs in complex odor environments.

An optogenetics approach aimed at identifying olfactory neurons which guide attraction or repulsion supports this hypothesis. The experiments by Bell and Wilson (28) involved using low-level optogenetic activation of OSNs in a two choice walking assay. The authors were able to obtain attractive and repulsive motor behavior upon stimulating eight OSN classes previously identified as attractive or repellent. The authors further examined the effects of stimulating two classes of OSNs on attractive or repulsive behaviors. These pairwise studies revealed that activation of certain OSNs resulted in behavioral output that summed linearly (were more attractive), but others did not. Pairing attractive OSNs with repellent OSNs did decrease attraction. Together, these results suggest that different OSNs can contribute different ‘weights’ towards an output behavior. The negative oviposition OSNs we identified in our study most likely add ‘negative’ weight to an olfactory-guided oviposition choice (56). Furthermore, it has been shown that certain glomeruli have greater influence over others (57). This observation gives rise to the possibility that a class of OSNs, in this case one mediating aversion, could act as a master switch and carry much more weight in the summation of antennal lobe inputs, giving that glomerulus the ability to ‘veto’ other inputs. It is unclear if this would be the case for the negative oviposition OSNs, but it is possible that OSNs that are sufficient to drive a specific behavior alone may carry more ‘weight’.

The olfactogenetic method can be used study an array of behaviors amenable to odor presentation. Ectopic expression of Or56a and activation by geosmin could be used to singly interrogate OSNs in many more olfactory contexts and allow for the widespread identification of putative receptors involved in behaviors including courtship, aggregation, or aggression. Furthermore, olfactogenetics may be a powerful tool used to interrogate how combinatorial OSN activities summate to produce any of these complex behaviors. With the stereotypic mapping of second order neurons, identifying primary inputs could lead to conclusions about higher order processing in olfactory cortex that ultimately elucidate how the brain evaluates and uses olfactory information to help animals survive.

## MATERIALS AND METHODS

***Fly Stocks:*** Wildtype flies were *IsoD1 (w^1118^)*, and all lines used in behavioral experiments including the two *Or56a* knockout lines were backcrossed for five generations to wild type. All *OrX-Gal4* lines were crossed into the outcrossed *Or56a* knockout background. *Gal4* lines used for this study are listed in Lin & Potter 2015, Table 1(23). Flies used for *Orco* mutant experiments contained two different alleles as reported in (29).

***Generation of the UAS-Or56a fly line:*** The *Or56a* coding region was PCR amplified from *IsoD1 (w^1118^)* genomic DNA using primers with 15 bp extensions appropriate for InFusion cloning (5’-GAATAGGGAATTGGGAATTCATGTTTAAAGTTAAGGATCTGTTGC-3’ and 5’-ATCTGTTAACGAATTCCTAATACAAGTGGGAGCTACG-3’). InFusion cloning (Clonetech Laboratories, Inc) was used to subclone *Or56a* into the EcoRI cut site in the multiple cloning region of *pUAST* (58). This vector was then injected into embryos for P-element insertion.

***Generation of the Or56a Knockout through accelerated homologous recombination:*** A deletion mutant was generated using accelerated gene targeting as reported in (59). Briefly, 4559 bps of genomic sequence immediately upstream and 3021 bps immediately downstream to *Or56a* were PCR amplified using primers designed for InFusion cloning to create a 5’ and 3’ homology arm, respectively (**Or56a_5’homarm_REV** 5’-AGTTGGGGCACTACG**GTTAAACTGTTTAGCGTTAACCATATTC**-3′, **Or56a 5’homarm FOR2** 5’-CTAGCACATATGCAGC**TCACAGCGCTTGTCGTAAT**-3′; **Or56a_3′homarm_FOR** 5-ACGAAGTTATCA**AGGGAAAGCCTTTTCTTCAGG**-3’, Or56a_3’homarm_REV2 5’ -GATCTTTACTAG**TTTTCCGCTTCTGCTCTACG**-3’;

Bolded nucleotides represent genomic sequence, and unbolded sequence indicates vector nucleotides. Sequentially, the 3’ homology arm was InFusion cloned into the *SpeI* restriction site of MCS B in *pTV*^*Cherry*^ (vector from lab of J.P. Vincent)’ and the 5’ homology was cloned into the *NheI* site of the MCS A. This knockout construct was used to generate a ‘Donor’ line that was crossed to *hs-Flp*, *hs-SceI* (BS#25679). Flies were heat-shocked at 37°C, 48 hours and 72 hours after egg-laying for 1 hour each. Female progeny of the heat shocked flies were then screened for mottled eyes and crossed with *ubi-Gal4[pax-GFP]* to select against off-target recombination events. The *Or56aKI* was validated with primers *GAL4-FOR* (5’- TCGATACCGTCGACTAAAGCC -3’) and *Or56a-TEST-REV2* (5’- AAAATCGAGGGGCTAAACAGTGTC -3’), as shown in Figure S1.

***Chemicals:*** Geosmin in methanol was purchased from Sigma-Aldrich at highest available purity, ≥97% (Product #: G5908-1ML, Lot #: BCBP7178V). Chemical as received was dried to remove methanol and then diluted to 4 mg/mL in mineral oil (Sigma-Aldrich, Product #: 330779-1L, Lot #: MKBF6530V). Methyl laurate (Product #: 234591-2.5G, Lot #: BCBQ6830V), (Z)-9-tricosene (Product #: 859885-1G, Lot #: 04706LDV) and farnesol (Product #: F203-25G, Lot #: MKBG0101V) were purchased from Sigma-Aldrich.

***SSR:*** All sensilla were identified using fluorescence from either *10X-UAS-IVS-mCD8-GFP* (II) or *15X-UAS-IVS-mCD8-GFP* (III) recombined onto *OrX-Gal4* lines. These recombined lines were crossed to *UAS-Or56a* to test the efficacy of misexpressing Or56a in non-ab4 neurons. Single sensillum recordings from the *OrX-Gal4’s* cognate sensillum were obtained using methods and lines described in (23).

***Oviposition Assay:*** Equal numbers of female and male adult flies were collected within 24 hours of eclosion and group housed in fly food vials for three days. On day four, all flies were transferred to a vial with only wet yeast paste to prime females for egg laying. 50 mL of 1% agarose in double distilled water was allowed to cool to precisely 65°C. 1 uL of odorant or vehicle was pipetted into the 50 mL of 1% agarose. This solution was dispensed into each well of a three-well spot plate (Corning, Product No 7223-34 -discontinued (20 drops per well); Replica three-well spot plate printed in porcelain with matte black finish through Shapeways (14 drops per well)) using a pipet-aid with a 10 mL serological pipette (Danville Scientific, Part No: P7134). Flies were anesthetized on ice for 3-5 minutes, and males were removed. ~10 female flies were tapped onto each spot plate and the lid of a 100x20 mm tissue culture dish (Corning Incorporated, Product No: 353003) was placed on top to cover the top of the assay. The lip on the spot plate allows room for the flies to walk on and between the three wells. All experiments were begun between 5:00-7:00 pm, and flies were incubated on the assay in a dark, humidified incubator at 25°C and 89-94% humidity for 22-23 hours, and the number of eggs on the agarose of each well was counted. Counts were normalized to the number of flies loaded into each assay (# of eggs in well/number of flies). We discarded experiments in which flies laid fewer than 8 eggs/fly/day.

Oviposition index was calculated as follows:

OI = (O - NO^avg^)/(O + NO^avg^)

OI = Oviposition Index
O = # of eggs in well containing odorant
NO^avg^ = Average # of flies between two no odorant vehicle control wells

***Statistics:*** Normality was determined using the Bartlett's Test of Homogeneity of Variances (p < 0.01). Given that the data meet requirements for running parametric tests, an ANOVA shows that at least two of the means from the experimental groups are different from one another (p = 2.76e-13). The post hoc Dunnett's Many-to-One Multiple Comparisons test, with each experimental group compared to the Or56a knockout control, indicates that 9 out of the 23 tested OrX-Gal4 lines statistically significantly induce aversive behavior in our oviposition assay. These tests were all run using the native and ‘multcomp’ statistics packages in R.

## ACKNOWLEDGEMENTS

We thank R. Benton, J. Carlson, and the Bloomington Drosophila Stock Center (NIH P40OD018537) for fly lines. We thank A. Kolodkin, M. Wu, and M. Caterina for discussions. We thank R. Du for assistance generating the *Or56a* knockout. This work was supported by the NIH NIDCD (R01DC013070, CJP).

## SUPPLEMENTAL METHODS

### Projection Neuron tracing and analysis (Figure S4)

Registered confocal images containing labeled single neurons were obtained from M. Costa and G. Jefferis (60) based on original raw data from the FlyCircuit dataset [NCHC (National Center for High-performance Computing) and NTHU (National Tsing Hua University), Hsinchu, Taiwan] (10). 3D reconstructions of single neurons were created with semi-automated tracing in Amira software (Zuse Institute, Berlin, Germany) using the hxskeletonize plugin (61). These 3D reconstructions were then pooled together with earlier published sets of traced neurons (9, 62, 63) producing a dataset of 129 neurons for all the PN types for which behavioral data was available.

The 3D *dot properties* representations (60, 64) for reconstructed neurons were calculated by using the nat packages created by G. Jefferis and J. Manton for the R statistical computing environment (https://jefferislab.github.io). Neurons on the right side of the brain were flipped over to the left side by mirroring and applying flipping registration to them using the https://github.com/jefferislab/nat.templatebrains and https://github.com/jefferislab/nat.flybrains R packages (65). The PN axonal projections in an area of interest including the lateral horn of the brain were separated from the rest of the neuron using the nat package. NBLAST was then used to perform all by all comparisons of these arbors; the raw NBLAST scores were normalized (mean of forward and reverse scores/sum of forward and reverse scores) and then aggregated per glomerulus by taking the arithmetic mean of normalized scores for each PN type. Finally, the resultant all-glomeruli-by-all-glomeruli NBLAST score matrix was converted into a distance matrix by subtracting the mean value from 1 for each PN type. This distance matrix was then used for hierarchical clustering and statistical testing.

To compare the anatomical similarity of the 5 behaviorally most significant PN types to the rest of the sample, random groups of 5 PN types were drawn from the sample without replacing (“jackknifing”) and the obtained empirical cumulative distribution function of distance scores was then compared to the observed distance scores of the negative-oviposition PNs.

To examine the relationship between behavioral OI values and PN morphology in the LH we also converted OI values to a distance matrix (using Euclidean distance) and used a Mantel test to look for a correlation between the two matrices. Secondly, we used the most significant negative oviposition PN type as a reference point and looked for a correlation between the anatomical distance to the most negative oviposition PN type and the raw OI values by a Pearson’s product moment correlation test. To avoid observing a false positive correlation, the most negative oviposition PN type was removed from the sample for this analysis, as it by definition would have both a low OI value and a distance score of 0.

All statistical testing was done using the R statistical computing environment.

**Figure S1:**
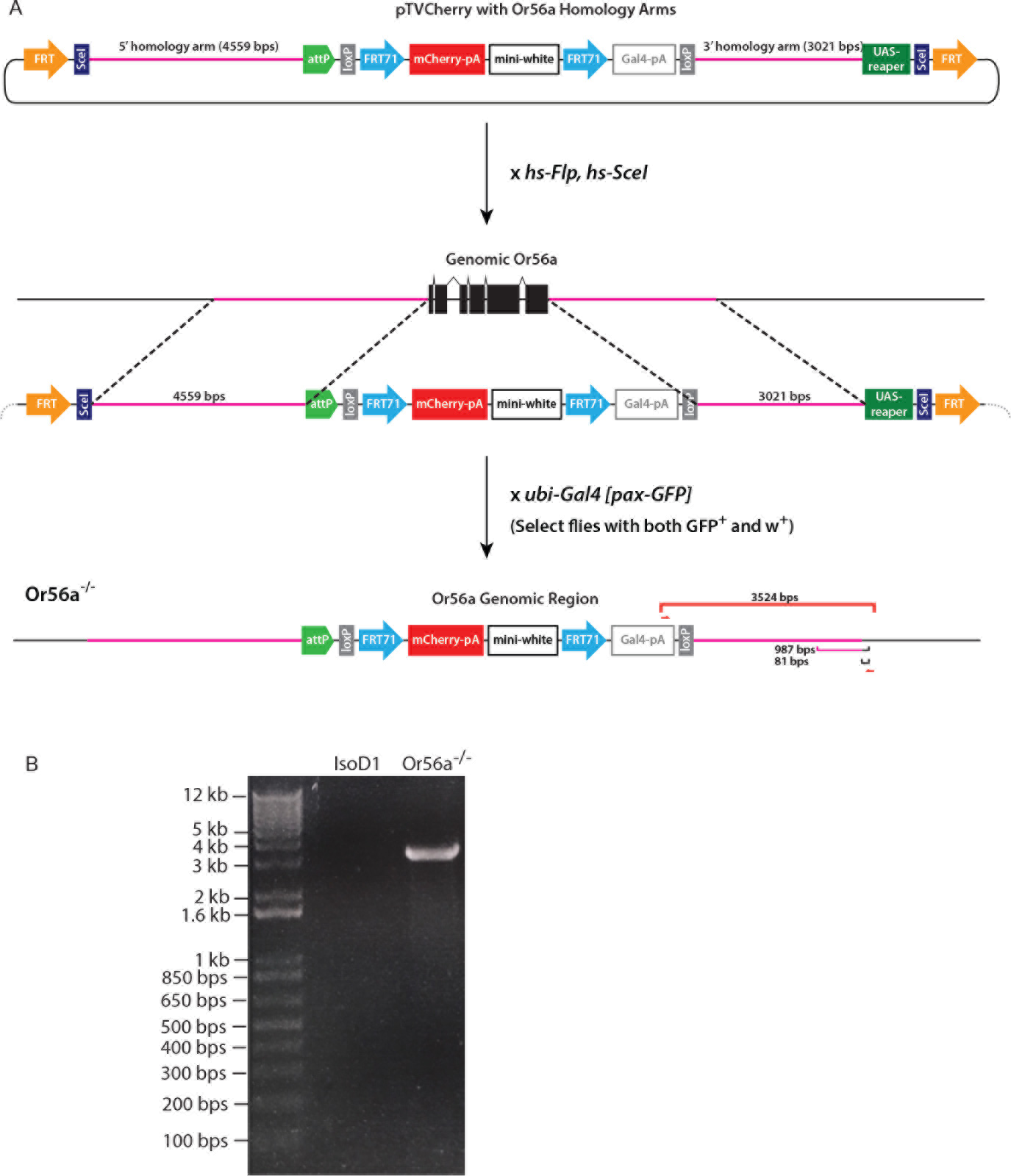
Generation of *Or56a*^-/-^ mutant. The accelerated homologous recombination method (59) was used to generate an *Or56a* knockout. **A)** Schematics of the cloning of *Or56a* homology arms into *pTV*^*Cherry*^, and the resulting *Or56a* mutant locus. The primer positions for validating correct insertion of knockin construct into Or56a locus of genome are also shown. **B)** PCR verification of the *Or56a* mutants using primers shown in A.

**Figure S2:**
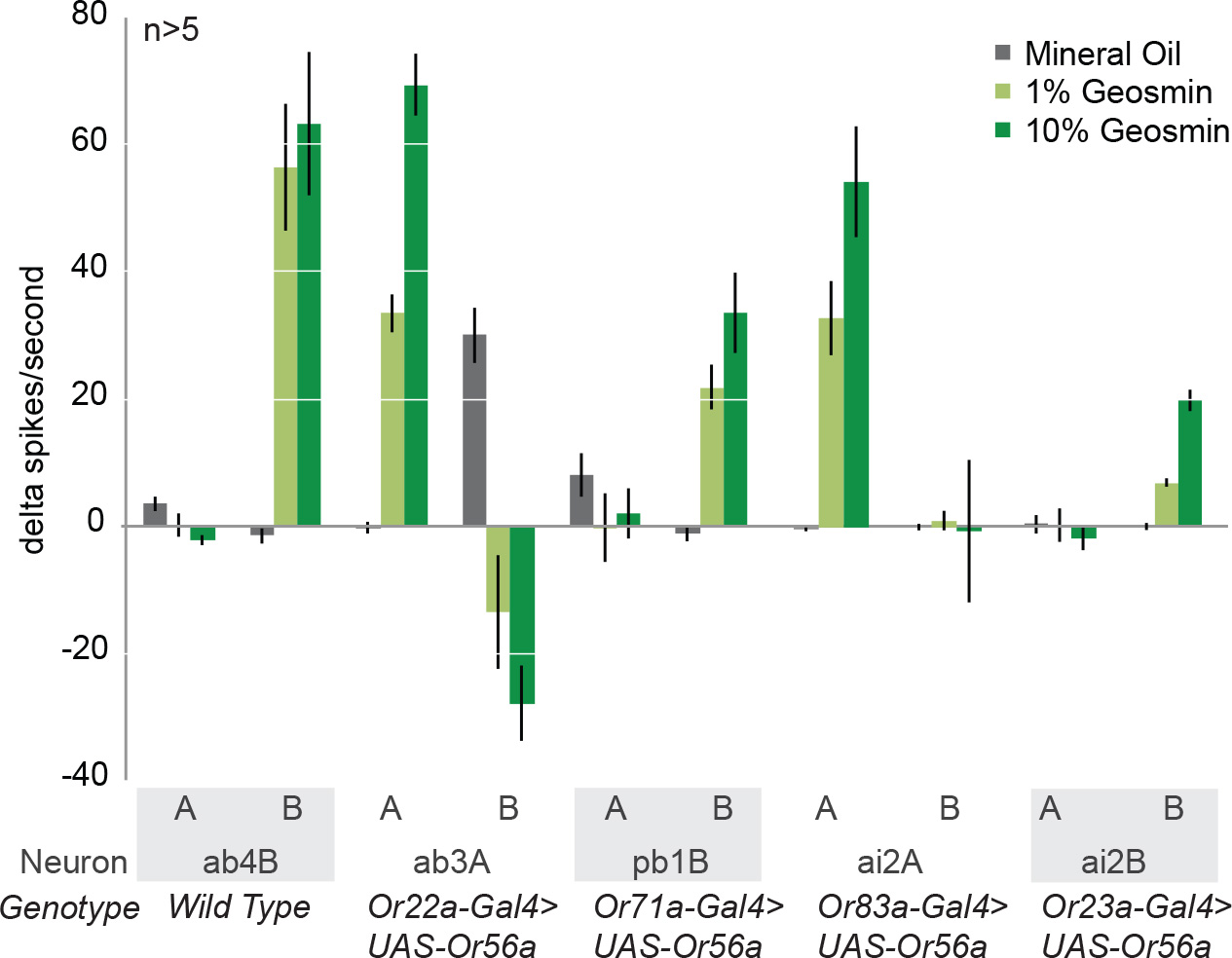
Responses of olfactory neurons in a sensilla containing a neuron expressing *Or56a*. Quantifications of all neurons within a targeted sensilla. ‘Neuron’ lists the olfactory neuron expressing *Or56a* in the sensilla. n>5 for all except ai2A (n=3). Error bars are SEM. Related to Figure 2.

**Figure S3:**
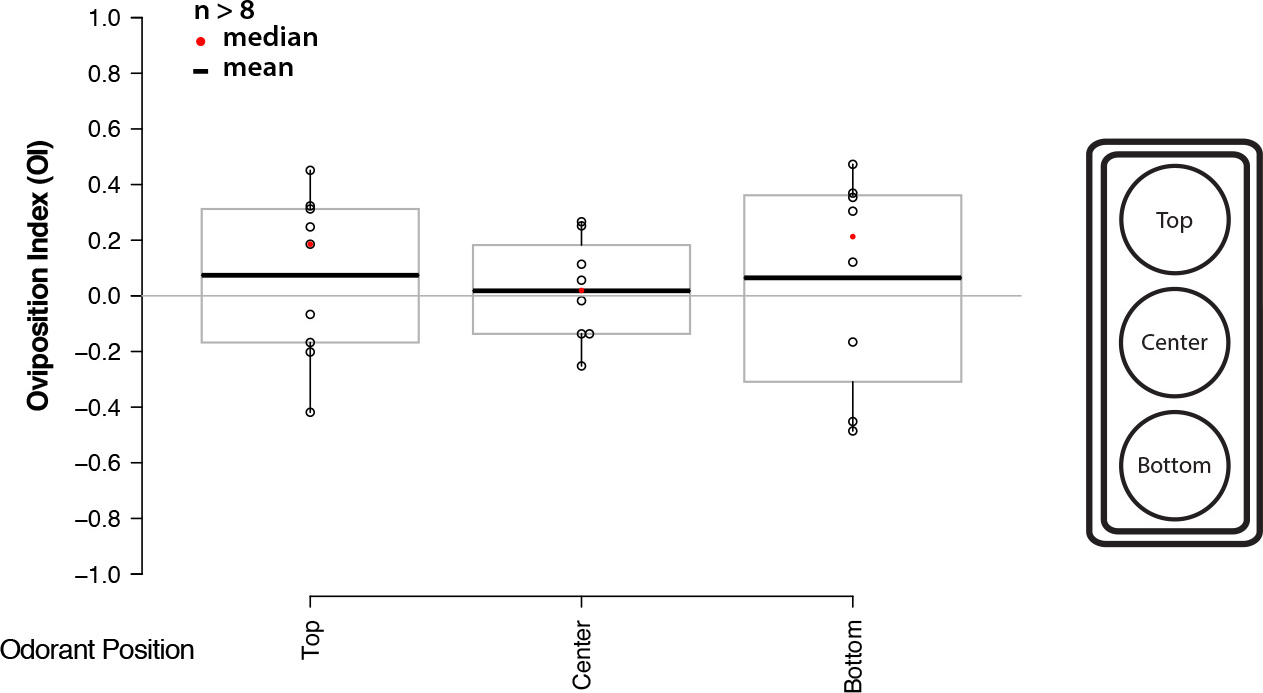
Positional Controls for Two-Choice Oviposition Assay. The position of the Odorant Well in the three-well assay does not affect behavior. Differences are not statistically significant as determined by a Dunnett's Many-to-One Comparisons Test.

**Figure S4:**
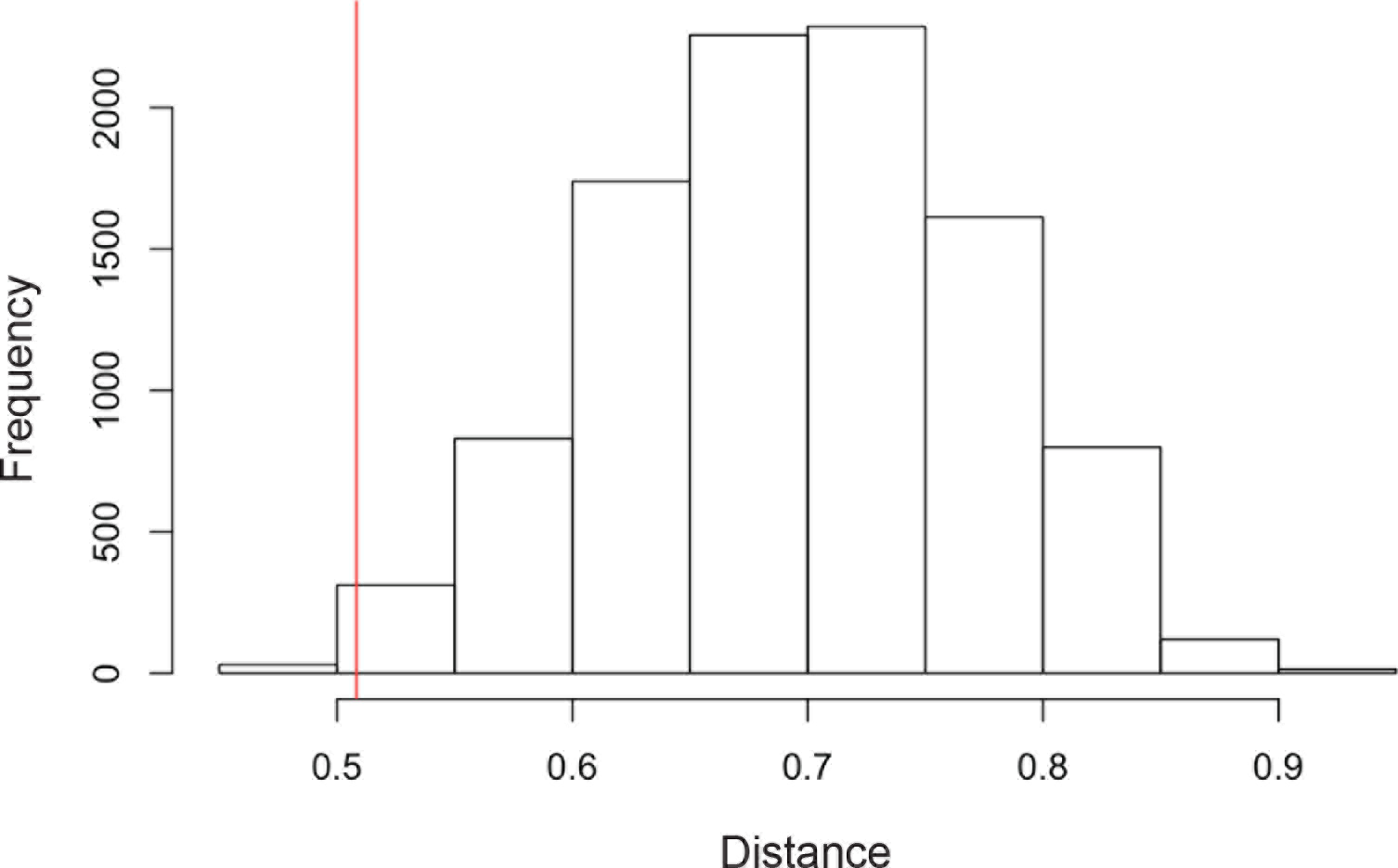
Oviposition neurons are statistically more similar to each other than to other known PNs. Histogram of morphological distances between lateral horn processes of pairs of neurons based on all known PN anatomical traces. Distances can range from 0 (identical) to 1.8 (maximally different). Negative oviposition neurons appear statistically more similar to one another (mean distance=0.51, red line) compared with the distribution for random pairs of neurons (grey bars, mean distance=0.69). p = 0.0035, based on 10,000 randomly sampled pairs.

**Table S1.** Olfactory receptors implicated in oviposition decisions.

| Olfactory Receptor | Chemical/Receptor Identification Method | Odorant | Citation |
| --- | --- | --- | --- |
| Or7a | Behavior, GCMS and SSR | 9-tricosone | Lin et al, 2015 |
| Or19a | GC-SSR screen/Ca2+ imaging | citrus volatiles | Dweck et al, 2013 |
| Or49a | GC-SSR screen/Ca2+ imaging | parasitoid wasp pheromones | Ebrahim et al, 2015 |
| Or56a | SSR screen | geosmin | Stensmyr et al, 2012 |
| Or67d | Behavior, GCMS and SSR | cVA | Bartelt et al, 1985 |
| Or71a | GC-SSR screen with natural odorant | dietary antioxidants | Dweck et al, 2014 |

**Table S2.** Bloomington Stock numbers for lines used in Oviposition screen.

| Gal4 Line | Bloomington Stock Number |
| --- | --- |
| Or7a | KI from Lin and Potter, 2015 |
| Or19a | 23887 |
| Or22a | 9951 |
| Or23a | 9956 |
| Or35a | 9967 |
| Or42a | 9969 |
| Or42b | 9971 |
| Or43a | 9974 |
| Or47a | 9982 |
| Or47b | 9983 |
| Or49a | 9985 |
| Or59c | 23899 |
| Gr63a | 9942 |
| Or65a | 9994 |
| Or67b | 9995 |
| Or67d | KI from Kurtovic, 2007 |
| Or71a | 23122 |
| Or82a | 23126 |
| Or83c | 23131 |
| Or85a | 24461 |
| Or85d | 24148 |
| Or88a | 23138 |
| Or92a | 23139 |

## REFERENCES

1. Masse NY, Turner GC, & Jefferis GSXE (2009) Olfactory information processing in Drosophila. Curr Biol 19(16):R700–713.

2. Kaupp UB (2010) Olfactory signalling in vertebrates and insects: differences and commonalities. Nat Rev Neurosci 11(3): 188–200.

3. Hallem EA & Carlson JR (2006) Coding of odors by a receptor repertoire. Cell 125(1): 143–160.

4. Mansourian S & Stensmyr MC (2015) The chemical ecology of the fly. Curr Opin Neurobiol 34:95–102.

5. de Bruyne M, Foster K, & Carlson JR (2001) Odor coding in the Drosophila antenna. Neuron 30(2):537–552.

6. Grabe V, et al. (2016) Elucidating the Neuronal Architecture of Olfactory Glomeruli in the Drosophila Antennal Lobe. Cell Reports 16(12):3401–3413.

7. Fishilevich E & Vosshall LB (2005) Genetic and functional subdivision of the Drosophila antennal lobe. Current Biology 15(17):1548–1553.

8. Couto A, Alenius M, & Dickson BJ (2005) Molecular, anatomical, and functional organization of the Drosophila olfactory system. Current Biology 15(17):1535–1547.

9. Jefferis GSXE, et al. (2007) Comprehensive maps of Drosophila higher olfactory centers: spatially segregated fruit and pheromone representation. Cell 128(6): 1187–1203.

10. Chiang A-S, et al. (2011) Three-dimensional reconstruction of brain-wide wiring networks in Drosophila at single-cell resolution. Curr Biol 21(1): 1–11.

11. Yang CH, Belawat P, Hafen E, Jan LY, & Jan YN (2008) Drosophila egg-laying site selection as a system to study simple decision-making processes. Science 319(5870): 1679–1683.

12. Lihoreau M, Poissonnier LA, Isabel G, & Dussutour A (2016) Drosophila females trade off good nutrition with high-quality oviposition sites when choosing foods. J Exp Biol 219(Pt 16):2514–2524.

13. Markow TA & O’Grady P (2008) Reproductive ecology of Drosophila. Functional Ecology 22(5):747–759.

14. Zhu EY, Guntur AR, He R, Stern U, & Yang CH (2014) Egg-laying demand induces aversion of UV light in Drosophila females. Curr Biol 24(23):2797–2804.

15. Lin C-C, Prokop-Prigge KA, Preti G, & Potter CJ (2015) Food odors trigger Drosophila males to deposit a pheromone that guides aggregation and female oviposition decisions. eLife 4:44.

16. Stensmyr MC, et al. (2012) A Conserved Dedicated Olfactory Circuit for Detecting Harmful Microbes in Drosophila. Cell 151(6):1345–1357.

17. Dweck HKM, et al. (2013) Olfactory Preference for Egg Laying on Citrus Substrates in Drosophila. Current Biology:1–9.

18. Dweck HKM (2014) Drosophila Olfactory Neuroecology-Function and Evolution. eLife 3:1–137.

19. Ebrahim SAM, et al. (2015) Drosophila Avoids Parasitoids by Sensing Their Semiochemicals via a Dedicated Olfactory Circuit. PLoS Biol 13(12):e1002318–1002318.

20. Bartelt RJ, Schaner AM, & Jackson LL (1985) cis-Vaccenyl acetate as an aggregation pheromone in Drosophila melanogaster. J Chem Ecol 11(12): 1747–1756.

21. Klapoetke NC, et al. (2014) Independent optical excitation of distinct neural populations. Nat Meth:1–14.

22. Hamada FN, et al. (2008) An internal thermal sensor controlling temperature preference in Drosophila. Nature 454(7201):217–220.

23. Lin C-C & Potter CJ (2015) Re-Classification of Drosophila melanogaster Trichoid and Intermediate Sensilla Using Fluorescence-Guided Single Sensillum Recording. PloS one 10(10):e0139675.

24. Laughlin JD, Ha TS, Jones DNM, & Smith DP (2008) Activation of pheromone-sensitive neurons is mediated by conformational activation of pheromone-binding protein. Cell 133(7):1255–1265.

25. Benton R, Vannice KS, & Vosshall LB (2007) An essential role for a CD36-related receptor in pheromone detection in Drosophila. Nature 450(7167):289–293.

26. Ronderos DS, Lin CC, Potter CJ, & Smith DP (2014) Farnesol-Detecting Olfactory Neurons in Drosophila. Journal of Neuroscience 34(11):3959–3968.

27. Su C-Y, Menuz K, Reisert J, & Carlson JR (2012) Non-synaptic inhibition between grouped neurons in an olfactory circuit. Nature:1–7.

28. Bell JS & Wilson RI (2016) Behavior Reveals Selective Summation and Max Pooling among Olfactory Processing Channels. Neuron:1–40.

29. Larsson MC, et al. (2004) Or83b encodes a broadly expressed odorant receptor essential for Drosophila olfaction. Neuron 43(5):703–714.

30. Kwon JY, Dahanukar A, Weiss LA, & Carlson JR (2007) The molecular basis of CO2 reception in Drosophila. Proc Natl Acad Sci USA 104(9):3574–3578.

31. Clyne PJ, Warr CG, & Carlson JR (2000) Candidate taste receptors in Drosophila. Science 287(5459):1830–1834.

32. Scott K, et al. (2001) A chemosensory gene family encoding candidate gustatory and olfactory receptors in Drosophila. Cell 104(5):661–673.

33. Sato K, et al. (2008) Insect olfactory receptors are heteromeric ligand-gated ion channels. Nature 452(7190):1002–1006.

34. Wicher D, et al. (2008) Drosophila odorant receptors are both ligand-gated and cyclic-nucleotide-activated cation channels. Nature 452(7190): 1007–1011.

35. Dweck HKM, et al. (2015) Pheromones mediating copulation and attraction in Drosophila. Proc Natl Acad Sci USA:201504527.

36. Lin H-H, et al. (2016) Hormonal Modulation of Pheromone Detection Enhances Male Courtship Success. Neuron 90(6): 1272–1285.

37. Vosshall LB (2007) Into the mind of a fly. Nature 450(7167): 193–197.

38. Wasserman S, Salomon A, & Frye MA (2013) Drosophila Tracks Carbon Dioxide in Flight. Curr Biol.

39. Lone SR, Venkataraman A, Srivastava M, Potdar S, & Sharma VK (2015) Or47b-neurons promote male-mating success in Drosophila. Biol Lett 11(5):20150292.

40. Schlief ML & Wilson RI (2007) Olfactory processing and behavior downstream from highly selective receptor neurons. Nat Neurosci 10(5):623–630.

41. Barlow HB (1961) Possible principles underlying the transformations of sensory messages. Sensory Communication, ed Rosenblith WA (MIT Press), pp 217–234.

42. Joseph RM & Heberlein U (2012) Tissue-specific Activation of a Single Gustatory Receptor Produces Opposing Behavioral Responses in Drosophila. Genetics 192(2):521–532.

43. Joseph RM, Devineni AV, King IF, & Heberlein U (2009) Oviposition preference for and positional avoidance of acetic acid provide a model for competing behavioral drives in Drosophila. Proc. Natl. Acad. Sci. U.S.A. 106(27): 11352–11357.

44. Dweck HKM, Ebrahim SAM, Farhan A, Hansson BS, & Stensmyr MC (2015) Olfactory Proxy Detection of Dietary Antioxidants in Drosophila. Current Biology.

45. Mattheis JP & Roberts RG (1992) Identification of geosmin as a volatile metabolite of Penicillium expansum. Appl Environ Microbiol 58(9):3170–3172.

46. Gerber NN & Lechevalier HA (1965) Geosmin, an earthly-smelling substance isolated from actinomycetes. Appl Microbiol 13(6):935–938.

47. Vertuani S, Angusti A, & Manfredini S (2004) The antioxidants and pro-antioxidants network: an overview. Curr Pharm Des 10(14):1677–1694.

48. Jimenez-Del-Rio M, Guzman-Martinez C, & Velez-Pardo C (2010) The effects of polyphenols on survival and locomotor activity in Drosophila melanogaster exposed to iron and paraquat. Neurochem Res 35(2):227–238.

49. Semmelhack JL & Wang JW (2009) Select Drosophila glomeruli mediate innate olfactory attraction and aversion. Nature 459(7244):218–223.

50. Riffell JA (2012) Olfactory ecology and the processing of complex mixtures. Curr Opin Neurobiol 22(2):236–242.

51. Montague SA, Mathew D, & Carlson JR (2011) Similar odorants elicit different behavioral and physiological responses, some supersustained. J Neurosci 31(21):7891–7899.

52. Andersson MN, Löfstedt C, & Newcomb RD (2015) Insect olfaction and the evolution of receptor tuning. Frontiers in Ecology and Evolution 3(53).

53. Suh GSB, et al. (2004) A single population of olfactory sensory neurons mediates an innate avoidance behaviour in Drosophila. Nature 431(7010):854–859.

54. Malnic B, Hirono J, Sato T, & Buck LB (1999) Combinatorial receptor codes for odors. Cell 96(5):713–723.

55. Wyatt TD (2014) Pheromones and animal behavior: chemical signals and signatures (Cambridge University Press).

56. Badel L, Ohta K, Tsuchimoto Y, & Kazama H (2016) Decoding of Context-Dependent Olfactory Behavior in Drosophila. Neuron 91(1): 155–167.

57. Fişek M & Wilson RI (2013) Stereotyped connectivity and computations in higher-order olfactory neurons. Nat Neurosci 17(2):280–288.

58. Brand AH & Perrimon N (1993) Targeted gene expression as a means of altering cell fates and generating dominant phenotypes. Development 118(2):401–415.

59. Baena-Lopez LA, Alexandre C, Mitchell A, Pasakarnis L, & Vincent JP (2013) Accelerated homologous recombination and subsequent genome modification in Drosophila. Development 140(23):4818–4825.

60. Costa M, Manton JD, Ostrovsky AD, Prohaska S, & Jefferis GS (2016) NBLAST: Rapid, Sensitive Comparison of Neuronal Structure and Construction of Neuron Family Databases. Neuron 91(2):293–311.

61. Evers JF, Schmitt S, Sibila M, & Duch C (2005) Progress in functional neuroanatomy: precise automatic geometric reconstruction of neuronal morphology from confocal image stacks. J Neurophysiol 93(4):2331–2342.

62. Grosjean Y, et al. (2011) An olfactory receptor for food-derived odours promotes male courtship in Drosophila. Nature.

63. Silbering AF, et al. (2011) Complementary Function and Integrated Wiring of the Evolutionarily Distinct Drosophila Olfactory Subsystems. J Neurosci 31(38): 13357–13375.

64. Masse NY, Cachero S, Ostrovsky AD, & Jefferis GS (2012) A mutual information approach to automate identification of neuronal clusters in Drosophila brain images. Front Neuroinform 6:21.

65. Manton JD, et al. (2014) Combining genome-scale Drosophila 3D neuroanatomical data by bridging template brains. bioRxiv.

